# Conserved mycobacterial sRNA B11 regulates lipooligosaccharide synthesis at post-transcriptional level

**DOI:** 10.1101/2024.05.30.596634

**Authors:** Chuan Wang, Cheng Bei, Yufeng Fan, Qingyun Liu, Yue Ding, Howard E Takiff, Qian Gao

## Abstract

Extractable glycolipids of mycobacteria, such as lipooligosaccharides (LOS), play key roles in responding to environmental stress and altering the host immune response. However, although the biosynthesis of LOS is likely controlled at multiple levels to ensure proper composition of the cell wall, the key regulators are currently unknown. Here, we studied B11, a conserved mycobacterial sRNA, and found that it post-transcriptionally regulates LOS synthesis in *Mycobacteria marinum*. Deletion of B11 alters the colony morphology and RNA sequencing combined with mass spectrometry identified several genes in the LOS synthesis locus that are regulated by B11. We found that B11 uses the cytosine-rich loops of its rho-independent transcriptional terminator to interact with guanine-tracks adjacent to the ribosome binding sites of its target genes, thereby impeding translation and promoting mRNA degradation by RNase E. These comprehensive functional studies of mycobacterial sRNA B11 demonstrate sRNA-based regulation of cell wall synthesis in mycobacteria.

**Importance:** Despite being identified for more than a decade, the functional characterization and regulatory mechanisms of mycobacterial sRNAs remain largely unexplored. We present here the most comprehensive functional study of mycobacterial sRNAs to date, employing convincible target screening using multifaceted experimental approaches and phenotype analysis. Our work reveals how synthesis of mycobacterial lipooligosaccharides (LOS), one of the crucial extractable glycolipids involved in environmental stress response and host immune modulation, is regulated at the post-transcriptional level by the conserved sRNA B11. Furthermore, our discovery of a highly conserved sRNA exhibiting distinct functions across mycobacterial species exemplifies divergent functional evolution among sRNAs.

## Introduction

Mycobacteria possess a cell envelope that plays an important role in host-pathogen interactions and susceptibility to antibiotics(1). The defining feature of the mycobacterial cell envelope is a complex cell wall outside of the typical bacterial plasma membrane, termed the mycomembrane, which is composed of noncovalently linked peptidoglycan, arabinogalactans and mycolic acids (2). The outer leaflet of the mycomembrane contains the extractable glycolipids, including such diverse elements as glyco-peptidolipids (GPLs), phenolic glycolipids (PGLs), trehalose mycolate (cord factor) and lipooligosaccharides (LOSs). Similar to the diverse O polysaccharides found in the lipopolysaccharides (LPS) of Gram-negative bacteria, the structures of mycobacterial glycolipids are species-specific. For example, more than 30 LOSs have been identified in different mycobacterial species, including *M.kansasii*, *M. canettii* and *M. marinum* (3), and all have a trehalose core attached to the varied lipidic moieties(4). The diverse mycobacterial glycolipids confer phenotypic differences and are also implicated in the pathogenicity of some mycobacteria. For example, the smooth-rough (S/R) colony morphotype variation is attributed to the presence or absence GPL in *M. abscessus* and LOS in *M. marinum*, *M. canettii* and *M. smegmatis* (5,6). The biosynthesis pathways of these glycolipids are well-defined and their synthesis has been shown to be controlled at the transcriptional level by regulatory factors such as Lsr2 and serine-threonine kinases (7,8), but exactly how this regulation responds to external and internal cues during growth and stasis remains unknown.

Post-transcriptional control by small regulatory RNAs (sRNAs) is a key element in the regulation of many biological processes, including cell wall synthesis(9). The majority of the currently known sRNAs are 50-250 nucleotides in length and typically regulate their target mRNAs through imperfect base pairing near the ribosome binding sites (RBS), thereby both inhibiting translation initiation and promoting mRNA degradation (10-12). Alternatively, base-pairing with sRNA outside of the RBS can promote translation of the target mRNAs by altering the secondary structure to facilitate ribosome binding to increase translation, or masking the RNase recognition motifs and decrease mRNA degradation (13,14). In addition to regulation at the post-transcriptional level, sRNA base-pairing can also modulate transcription by suppressing premature Rho-dependent transcription termination(15).

Despite sharing similar base-pairing mechanisms, sRNA-mediated regulation varies among different bacterial species. In Gram-negative bacteria, including *Escherichia coli* and *Salmonella*, regulation of most sRNAs is dependent on RNA chaperons Hfq or ProQ, which can accelerate sRNA-mRNA annealing and protect sRNAs from RNase E degradation (16,17). In Gram-positive bacteria, however, the role of Hfq and ProQ’s homologues in sRNA-mediated regulation is unclear. In low GC-content Gram-positive bacteria, such as *Bacillus subtilis* and *Staphylococcus aureus*, the deletion of Hfq had no impact on sRNA-mediated regulation(18), and the genomes of high GC-content bacteria, such as mycobacteria, lack Hfq or ProQ homologues. It is possible that the strength of base-stacking interactions with GC pairs makes the RNA chaperons unnecessary for sRNA-mRNA annealing, or alternatively, the high GC-content bacteria may have distinct RNA chaperones that have yet to be identified. Although several functional sRNAs have been described in different mycobacterial species, including *M. tuberculosis*(19-25), studies to define their biological function and regulatory mechanisms are scarce.

Here, we explored the biological function and regulatory mechanisms of sRNA B11 in *M. marinum*. By using different methods to discover B11-targets, we found that B11 controls several key genes in the LOS biosynthesis locus, which is distinct from the regulatory roles reported for B11 orthologues in *M. smegmatis* and *M. abscessus* (26,27). Furthermore, we found that *M. marinum* strains lacking B11 display a clear colony phenotype, suggesting that B11 acts as a negative posttranscriptional regulator of LOS synthesis. The work described here is, to our knowledge, the most comprehensive study of any mycobacterial sRNA and illustrates the involvement of RNA-based gene regulation in the synthesis of mycobacterial cell wall components.

## Results

### B11 is an abundant and stable sRNA in *M. marinum*

B11 is a 93 nt sRNA that was first identified in the *Rv3660-Rv3661* intergenic region of *M. tuberculosis* (19). The nucleotide sequence of B11 is more than 90% conserved in mycobacteria (Figure 1A), including a −10 motif (TATAGT) that matches the consensus sequences of sigma factor A promoters typically found upstream of housekeeping genes. Because studies in *M. tuberculosis* require strict biosafety infrastructure, we chose to work with the more tractable *Mycobacterium marinum* as a laboratory model to explore the biological function of B11. Transcriptomic analyses in different mycobacteria have shown that B11 is one the most abundant sRNAs (21,22,28). To confirm whether this was also true in *M. marinum,* we evaluated the expression of B11 during different growth phases in 7H9 rich medium using northern blots with *in vitro* synthesized B11 transcripts as standards. We found that B11 expression was low in lag phase at OD_600_∼0.3, but increased during the early exponential phase (OD_600_∼0.7) and remained high for at least 5 days while the cultures had reached an OD_600_ of 6.0 (Figure 1B). Intriguingly, we found that 5S RNA, whose presence is often used as a reference in northern blots of many bacteria, exhibited an expression pattern similar to B11, with low expression at lag phase and then steady levels beginning with exponential phase growth. Quantification with *in vitro* generated RNA suggested that B11 accounts for ∼0.1 % of total RNA by weight (∼5 ng of 5000 ng) at stationary phase (OD_600_∼6.0), corresponding to approximately 150-600 copies/cell. When compared with sRNAs such as SdsR in *E. coli* and *Salmonella*, which is present in ∼300 copies/cell, or RaiZ, present in ∼50 copies/cell(17,29), B11 appears to be relatively abundant in *M. marinum*. We also determined the stability of B11 by arresting transcription initiation with rifampicin and found that the half-life of B11 was more than 20 min (Figure 1C). When compared to the average 9.5 min half-life for *M. tuberculosis* mRNA and 5.2 min for *M. smegmatis* mRNA(30), B11 appeared to be quite stable. We also found that degradation of B11 resulted in a processed isoform that was resistant to further decay, and this isoform was observed during standard growth (Figure 1B). The high abundance and prolonged stability of B11 suggested that it could play an important role in the biology of mycobacteria.

**Figure 1.**
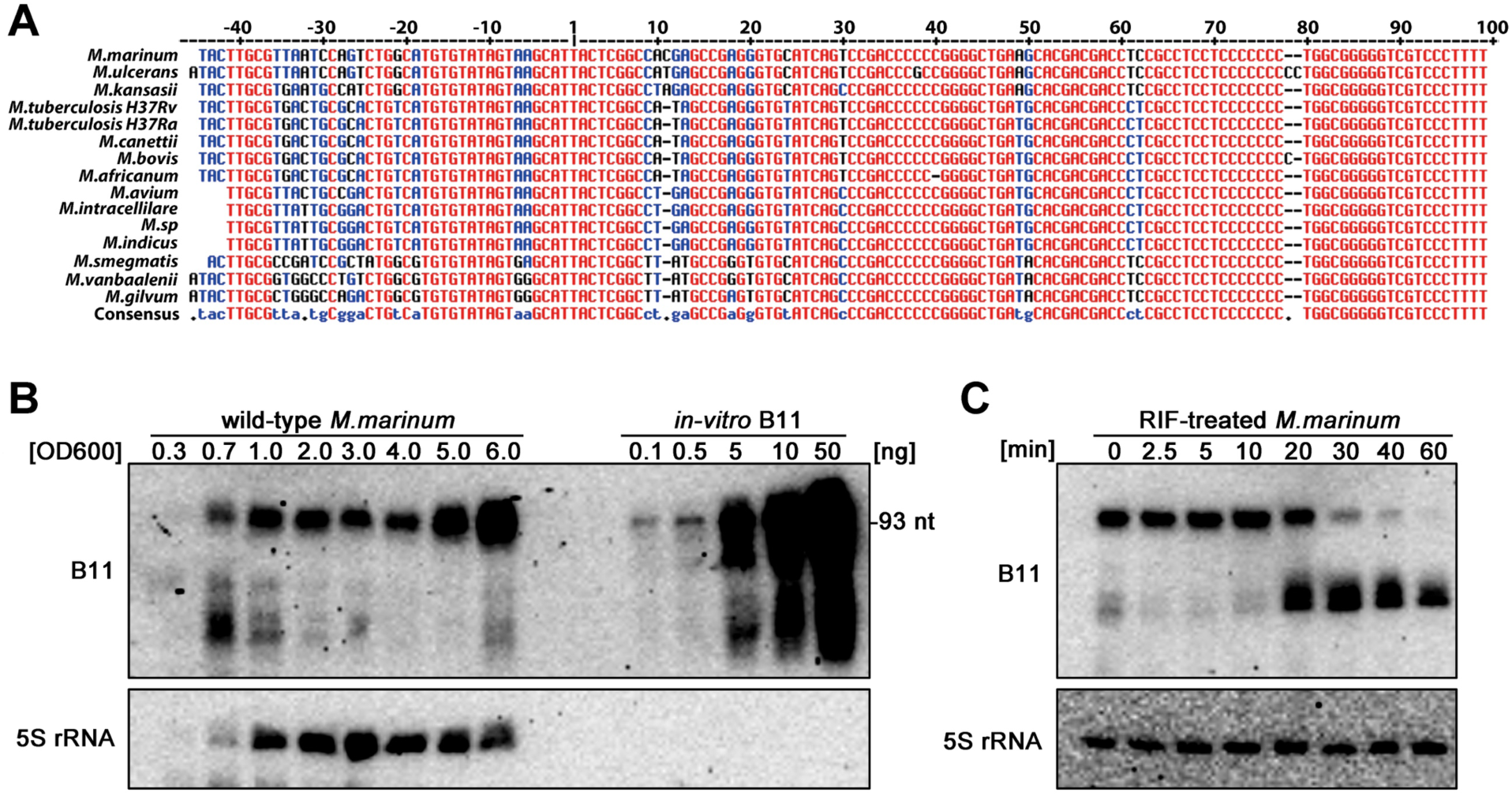
Characterization of sRNA B11 in mycobacterium. (A) Non-redundant alignment of the *b11* genes, including the upstream promoter regions, in different mycobacteria. All nucleotides are colored according to their degree of conservation: red, high conservation; blue, partial conservation; black, little or no conservation. ‘1’ marks the transcriptional start site. (B) Expression levels of B11 in an *M. marinum* wild-type strain through standard 7H9-OADC growth. Northern blot analysis of total RNA isolated from wild-type *M. marinum* grown to the indicated OD_600_. *In vitro* transcribed B11 was loaded as a standard to calculate the amount of *in vivo* B11. 5S RNA was used as loading controls. (C) Stability of B11. Wild type *M. marinum* strain was grown to OD _600_∼1.0 and treated with rifampicin at a final concentration of 200 μg/ml. Samples were collected at indicated time points and analyzed by northern blotting.

### Deletion of B11 altered *M. marinum* colony morphology

To identify the function of B11 in *M. marinum*, we constructed a B11-deleted strain (ΔB11) by replacing nucleotides 36-91 of B11 in the *M. marinum* genome with a kanamycin-resistance cassette, using a modified allelic exchange method based on temperature sensitive plasmid pPR27 (31). The deletion was confirmed by northern blots showing no expression of B11 in the deleted strain (ΔB11) during standard bacterial growth (Figure 2A). A slight growth defect was observed for the ΔB11 strain (ΔB11 + pCtr) during exponential phase growth in 7H9-OADC media, which was reversed by complementation of B11 expressed from its native promoter in multi-copy plasmid pSMT3 (ΔB11 + pP*_b11_*-B11) (Figure 2B). Intriguingly, the ΔB11 strain also displayed an altered colony morphology on 7H10 agar plates, with a reduction in the broad translucent border halo characteristic of wild-type *M. marinum* colonies (Figure 2C). Complementation of ΔB11 with a plasmid carrying intact B11 fully restored the wild-type colony morphology. As changes in mycobacterial colony morphology usually reflect alterations in the composition of cell wall glycolipids, the altered morphology of the ΔB11 strain suggested that B11 was involved with cell wall synthesis in *M. marinum*.

**Figure 2.**
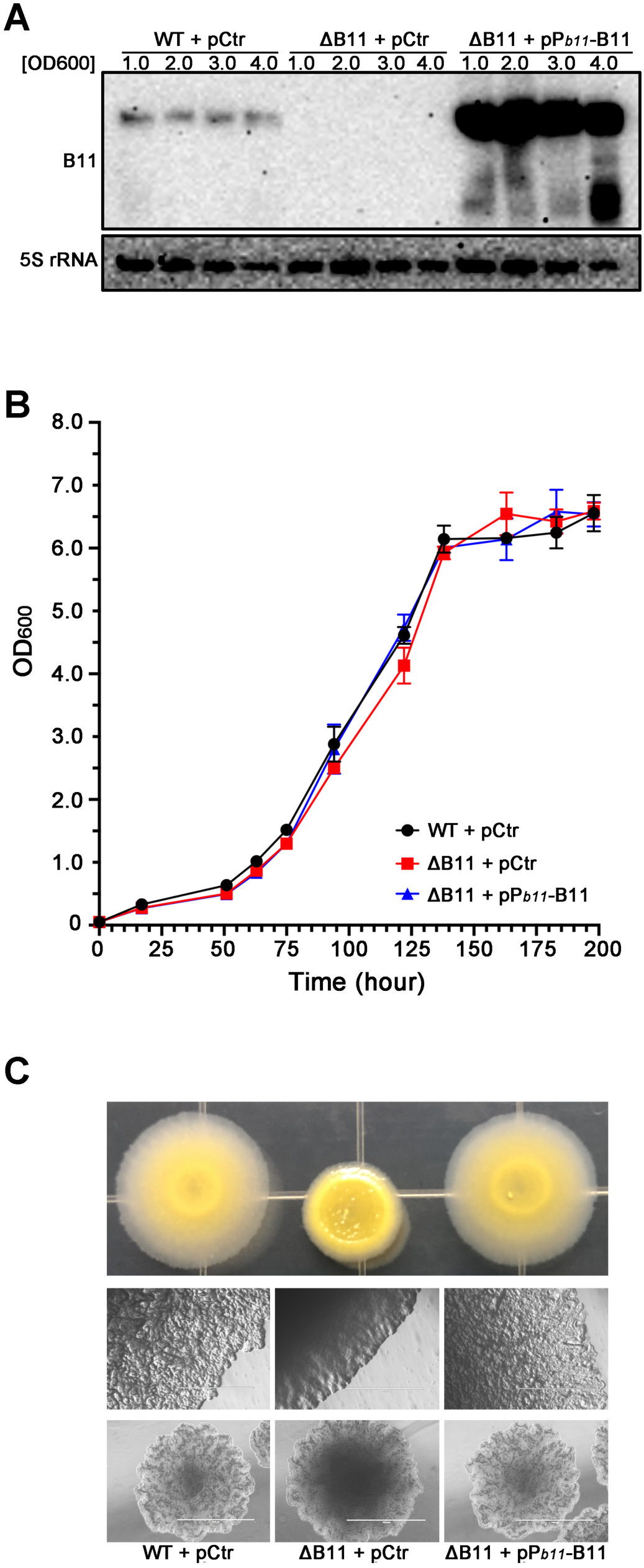
Deletion of B11 altered *M.marinum* colony morphology. (A) Expression levels of B11 in *M. marinum* wild-type (WT + pCtr), B11-deleted (ΔB11 + pCtr) and complemented (ΔB11 + pP*_b11_*-B11) strains. Northern blot analysis of total RNA isolated from different strains grown to the indicated OD _600_. 5S RNA was used as loading controls. (B) Growth curve of indicated *M.marinum* strains in 7H9-OADC. Error bars indicate standard deviations (n = 3). (C) Colony morphology of the indicated *M. marinum* strains in 7H10 agar. Strains as indicated above were grown to OD_600_ ∼1.0 in 7H9-OADC and then serially diluted and plated on 7H10-OADC agar, incubated for 5 days and visualized by camera (upper) or microscopy (middle). Single colonies of the indicated strains were also checked by microscopy (lower). Scale bar is 1000 μm.

### B11 represses the expression of genes from the lipooligosaccharides biosynthetic locus

To explore the molecular mechanism by which B11 regulates cell wall composition, we performed RNA sequencing and mass spectrometry to analyze differences in the transcriptomes and proteomes between a wild-type (WT + pCtr) strain, a B11-deleted (ΔB11 + pCtr) strain, and a strain with the B11-deletion that was complemented with over-expressed B11 (ΔB11 + pP*_b11_*-B11). RNA and protein samples were taken during early exponential phase (OD∼1.0), when the level of B11 expression in WT strains is stable. To identify B11-regulated genes, we performed two rounds of screening with different datasets (Figure 3A). The first-round sought genes whose expression differed between B11-deleted strains and strains complemented with overexpressed B11. We found 360 genes that surpassed the difference threshold (*p*<0.05) in both the RNA sequencing and mass spectrometry datasets (Figure 3B, Table S4), and the degree of the differences in the mRNA and protein levels were significantly correlated (*p*<0.0001, *r*^2^=0.3378, Simple linear regression). We then focused on the 35 genes with greater than 2-fold differences between B11-deleted and complemented strains in both mRNA and protein datasets. A second round of selection on these 35 genes compared their mRNA and protein levels in the wild-type versus B11-deleted strains and identified 4 candidates with significant differences (*p*<0.05): *mmar_1919*; *mmar_2909*; *mmar_2329*; and *mmar_4171*. Lastly, qRT-PCR on RNA samples prepared from five additional biological replicates (Figure 3C) confirmed that mRNA expression of three of the four candidate genes increased in the B11 deleted strain compared to the complemented strain. The expression difference could not be confirmed for *mmar_1919*.

**Figure 3.**
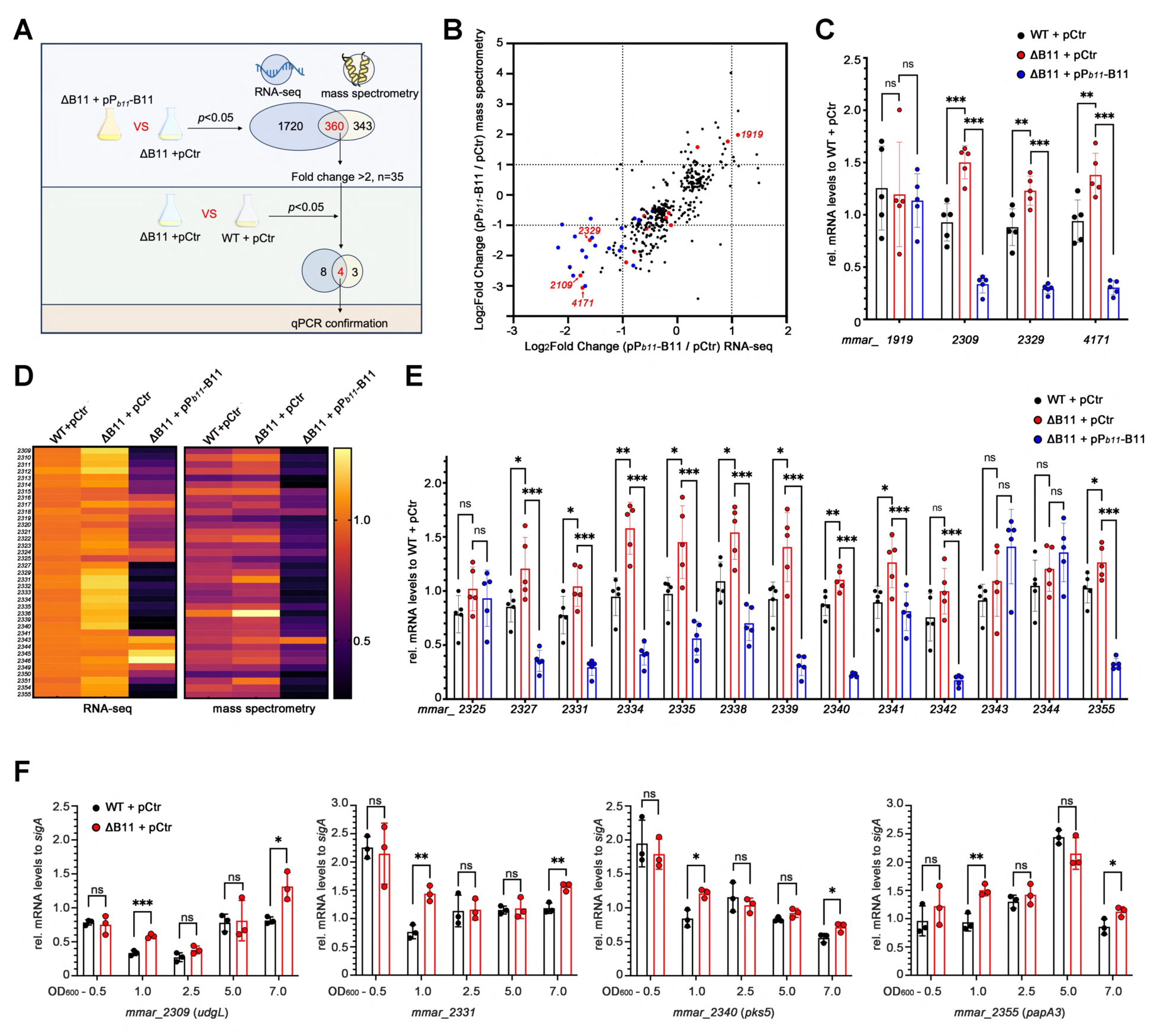
B11 represses the expression of genes from lipooligosaccharides biosynthetic locus. (A) Schematic overview of the workflow for transcriptomic and proteomic analysis. Total RNA and protein from indicated strains grown to OD _600_ ∼1.0 were analyzed by RNA sequencing and mass spectrometry. (B) Volcano plot displaying the log2 fold changes for genes with significant changes (*p*<0.05) in both the RNA sequencing (x-axis) and mass spectrometry data (y-axis) between B11-deleted (ΔB11 + pCtr) and complemented (ΔB11 + pP*_b11_*-B11) strains. 16 genes displaying significant gene expression changes (*p*<0.05) between wild-type and B11-deleted strains in both RNA sequencing and mass spectrometry are marked in red. Genes from the LOS biosynthetic locus (*mmar_2309-2355*) are marked in blue. Dashed lines refer to 2-fold changes. (C) Expression of 4 screened genes in the indicated strains confirmed by qRT-PCR. *sigA* was set as the reference gene for data analysis, and expression levels were normalized to one sample from the WT + pCtr group. Error bars indicate standard deviations (n = 5). (D) Heatmap displaying relative RNA and protein levels for genes from the *M. marinum* LOS locus (*mmar_2309-2355*), detected by RNA sequencing (left) and mass spectrometry (right). Expression levels were normalized to one sample from the WT + pCtr strain. Genes with undetected values were filtered. (E) Expression of selected genes from the LOS locus in the indicated strains measured by qRT-PCR and normalized to one sample from the WT + pCtr group. Error bars indicate standard deviations (n = 5). (F) Regulation of selected genes by B11 during different growth phases (n = 5). * *p*<0.05, ** *p*<0.01, *** *p*<0.001 and ns, no significant difference in a two-tailed *t* test.

We noticed that 2 of the 3 screened candidates, *mmar_2309*, mmar*_2329*, are located in the *mmar_2309-mmar_2346* genomic locus that spans approximately 60-kb (Figure S1) and is found only in *M. marinum.* This locus contains 38 genes encoding enzymes involved in the synthesis of lipooligosaccharides (LOS), which have been associated with the colony morphology of other mycobacteria species(32). Of note, both RNA sequencing and mass spectrometry revealed that nearly half of the genes in this locus were down-regulated when B11 was over-expressed (Figure 3D, Table S5), as were other genes close to this locus, *mmar_2350-2355*, which are also involved with LOS biosynthesis. Nine of the LOS associated genes had higher mRNA levels in the ΔB11 strain compared with wild-type strains, as shown by RNA sequencing and qRT-PCR on additional biological replicates (Figure 3D and 3E). Moreover, the differences in the expression of these genes between the ΔB11 and wild-type strains were seen in the early exponential (OD_600_∼1.0) and stationary phases (OD_600_∼7.0), but no differences were observed during mid (OD_600_∼2.5) and late exponential phases (OD_600_∼5.0), indicating that B11 regulation is growth phase dependent (Figure 3F). Taken together, these results suggest that B11 regulates the biosynthesis of lipooligosaccharides by targeting multiple genes in an *M. marinum*-specific genomic locus.

### Identification of B11-targets by *in vivo* MS2 affinity purification

Bacterial sRNAs commonly mediate post-transcriptional regulation through base pairing with the target mRNA(33,34). The presence of multiple B11-regulated genes in the LOS biosynthetic locus suggested that B11 may not regulate each of these genes individually, but may target only genes encoding enzymes upstream in the LOS biosynthesis pathway. To confirm the genes that B11 regulates directly through sRNA-mRNA binding, we used MS2 affinity purification to capture sRNA-mRNA interactions *in vivo*(35,36). In this approach, an MS2 RNA aptamer was attached to the 5’ end of B11 expressed from the strong Hsp60 promoter on plasmid pSMT3. mRNAs that interact with MS2-B11 RNA will be affinity purified from total RNA by binding to the MS2 coat protein (Figure 4A). MS2-B11 was successfully expressed as a 146 nt fragment (51 nt MS2, 2 nt UU linker and 93 nt B11), accompanied by additional processed fragments of sizes similar to untagged B11 (93 nt) (Figure 4B). The MS2-B11 RNA complemented the ΔB11 strain to restore the wild-type colony morphology, demonstrating that the B11 sequence remained functional (Figure 4C). A northern blot showed that the quantity of the MS2-B11 construct was reduced in the affinity flow-through and recovered in the elution sample (Figure 4D), indicating successful capture by affinity purification. By contrast, the level of untagged B11 was similar in the original lysate and the affinity flow-through and undetected in the elution sample. To confirm target enrichment with the MS2-B11 affinity purification, we selected 11 of the qRT-PCR validated gene targets in the LOS biosynthetic locus (Figure 3C and 3E) to check their mRNA abundance in the elution samples of cells expressing MS2-tagged or untagged B11. In addition, we selected the 10 genes encoding abundant proteins that mass spectrometry showed to have the greatest difference in protein levels when B11 was over-expressed (Figure S2), to evaluated which methods is more effective to capture the direct binding targets. Only 2 of these ten (*mmar_2309* and *mmar_4171*) showed similar B11 regulation in the RNA sequencing data. To minimize bias from systematic experimental variation, we used three sets of reference genes to normalize the qRT-PCR results: three genes in the LOS biosynthesis locus that are not regulated by B11 (*mmar _2325*, *mmar _2343* and *mmar _2344*); three genes randomly chosen from previous unpublished work in our laboratory (*mmar_3700*, *mmar_1863* and *mmar_4219*); and four genes commonly used as references for qRT-PCR (three rRNAs and *sigA*). Simple linear regression analysis with the *Ct* values of all 29 tested genes showed a similar distribution in the MS2-B11 and B11 samples (slope= 0.9202±0.025 and *r^2^* = 0.9003, Figure 4E). We then used quantile normalization to replace each *Ct* values with the average of that quantile across all tested genes(37). The degree of enrichment obtained by affinity purification was calculated by comparing the normalized *Ct* values in the eluted samples from the ΔB11 strains expressing either untagged B11 or tagged MS2-B11. The fold changes (MS2-B11 / B11) for the normalized *Ct* median values were 1.085 (median value) for rRNAs, 0.602 for the randomly chosen genes, 0.607 for non-B11 regulated genes, 0.520 for mass spectrometry screened genes, and 2.898 for the B11-regulated LOS synthesis genes, which were significantly enriched (*p*<0.05) when compared to any of the other gene groups (Figure 4F).

**Figure 4.**
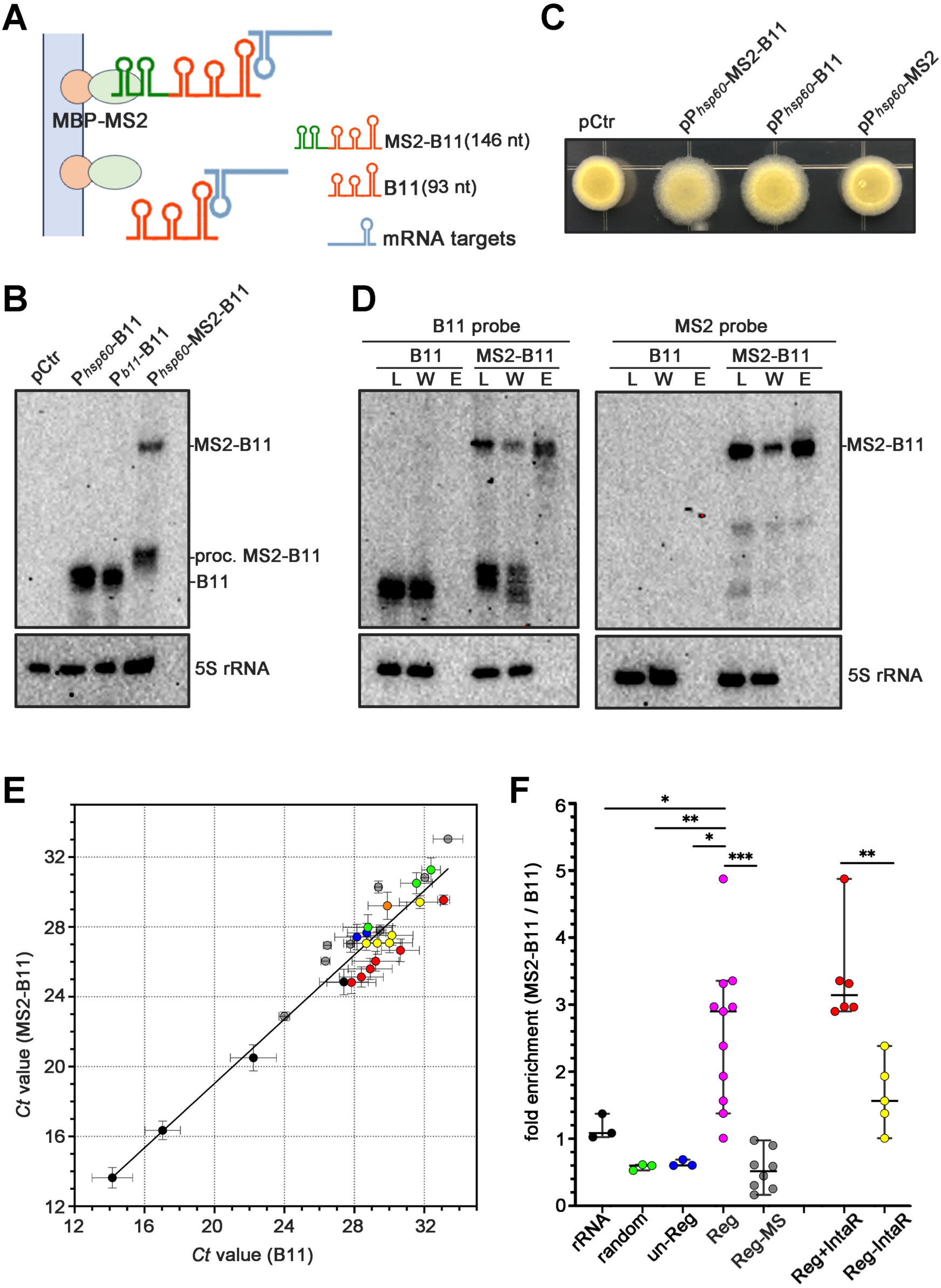
Identification of B11-targtes by *in vivo* MS2 affinity purification. (A) Schematic illustrating affinity purification of MS2 aptamer-tagged B11 to identify mRNA targets. The MS2 sequence was fused to the 5’ end of B11 and expressed *in vivo* from a plasmid in a B11-deleted strain of *M. marinum*. MS2-B11 RNA retains its capability for mRNA binding and can be captured by MBP-MS2 protein complex. (B) Northern blot analysis of total RNA isolated from B11-deleted strain carrying indicated plasmids. 5S RNA was used as loading controls. (C) Morphology of B11-deleted strain carrying indicated plasmids in 7H10 agar. (D) Northern blot analysis of RNA isolated from crude lysate (L), lysate washed through MBP column (W) and eluted samples (E) from B11-deleted strain carrying either P*_hsp60_*-MS2-B11 or P*_b11_*-B11 plasmids, and probed for B11 (Left) or MS2 (right). 5S RNA was used as loading controls. (E) Correlation of qRT-PCR *Ct* values of selected genes in RNA from elution samples of B11 deleted strains of *M. marinum* that contain plasmids with either B11 or MS2-B11. The dots indicate: genes that are regulated by B11 either with (red dots, Reg+IntaR, n=6) or without (yellow dots, Reg-IntaR, n=5) IntaRNA predicted binding regions; genes from the LOS biosynthesis locus that are not regulated by B11 (blue dots, un-Reg, n=3); randomly chosen genes from previous work in our laboratory (green dots, random, n=3); regulated proteins screened by mass spectrometry (gray dots, Reg-MS, n=8); and rRNAs (black dots, n=3). Simple linear regression analysis was performed with *Ct* values of all tested genes and the regression line was shown. (F) Fold enrichment after affinity purification for genes in the different groups, calculated after normalization. The dots from the indicated groups are marked with the same colors as in panel E, with the addition of B11-regulated LOS synthesis genes (collection of group Reg+ IntaR and Reg-IntaR), which are represented by magenta dots (Reg, n=11). Each group is represented by a marked bar indicating the median value along with a 95% confidence interval. * *p*<0.05, ** *p*<0.01, *** *p*<0.001, in a two-tailed *t* test.

We then used IntaRNA to predict the putative B11 binding sites in the 11 screened targets from the LOS synthesis locus. The five genes (*mmar _2330*, *mmar _2334*, *mmar _2335*, *mmar _2338, mmar _2341*) predicted to lack B11 binding sites had a median enrichment ratio of 1.565, whereas the 6 genes predicted to contain B11 binding sites (*mmar_2309*, *mmar_2331*, *mmar_2355* and *mmar_2340*-*2339 operon*, Figure S3) had a significantly higher median enrichment ratio of 3.139 (Figure 4F). This difference in fold enrichment suggests that B11 regulates the expression of these 6 targets directly by classical sRNA base-pairing *in vivo*. In addition, the median enrichment for the abundant proteins shown by mass spectrometry to have the greatest differences upon B11 over-expression, was only 0.520, and none of the genes were predicted to have B11 binding sites, indicating that they are not directly regulated by B11.

### Cysteine-rich loop in the transcriptional terminator of B11 is crucial for regulation

To explore the mechanistic details of B11-mediated repression, and confirm the region for base-pairing, the secondary structure of B11 was determined using *in vitro* structure-probing with single-strand-specific Pb (II). This analysis showed that B11 is highly structured with 3 intra stem-loops (Figure 5A and 5B), two of which are cytosine-rich: six cytosines residues in loop-2 and nine in loop-3. In addition, loop-3 has the structure of a typical rho-independent transcriptional terminator, with a GC-rich hairpin followed by a thymine rich segment (38). IntaRNA predicted that all 5 confirmed B11 gene targets harbor guanine-tracks near their ribosome biding sites, presumably allowing interaction with B11 cytosine-rich loops-2 or 3 (Figure S3). To confirm that base-pairing in these loops is involved in B11-mediated regulation, we replaced the cytosine residues in loops-2 and 3 with uridines. While replacement of the cytosines in loop-2 had no impact on B11 expression, replacing uridines for the cytosines in loop-3 significantly reduced the presence of B11 (Figure 5C), perhaps by affecting transcriptional termination. Furthermore, when the cytosine residues in loop-2 were replaced with uridines, there was no change in either cell morphology (Figure 5D) or mRNAs levels of the target genes compared to B11 with wild-type sequence (Figure 5E). By contrast, replacement of the cytosines in loop-3 abolished B11 mediated repression of target gene mRNA levels. Moreover, when the cytosines in both loop-2 and loop-3 were replaced with uridines, the effects were the same as when only loop-3 is mutated, suggesting that the C-track in B11 loop-3 is the critical region for B11-mediated regulation.

**Figure 5.**
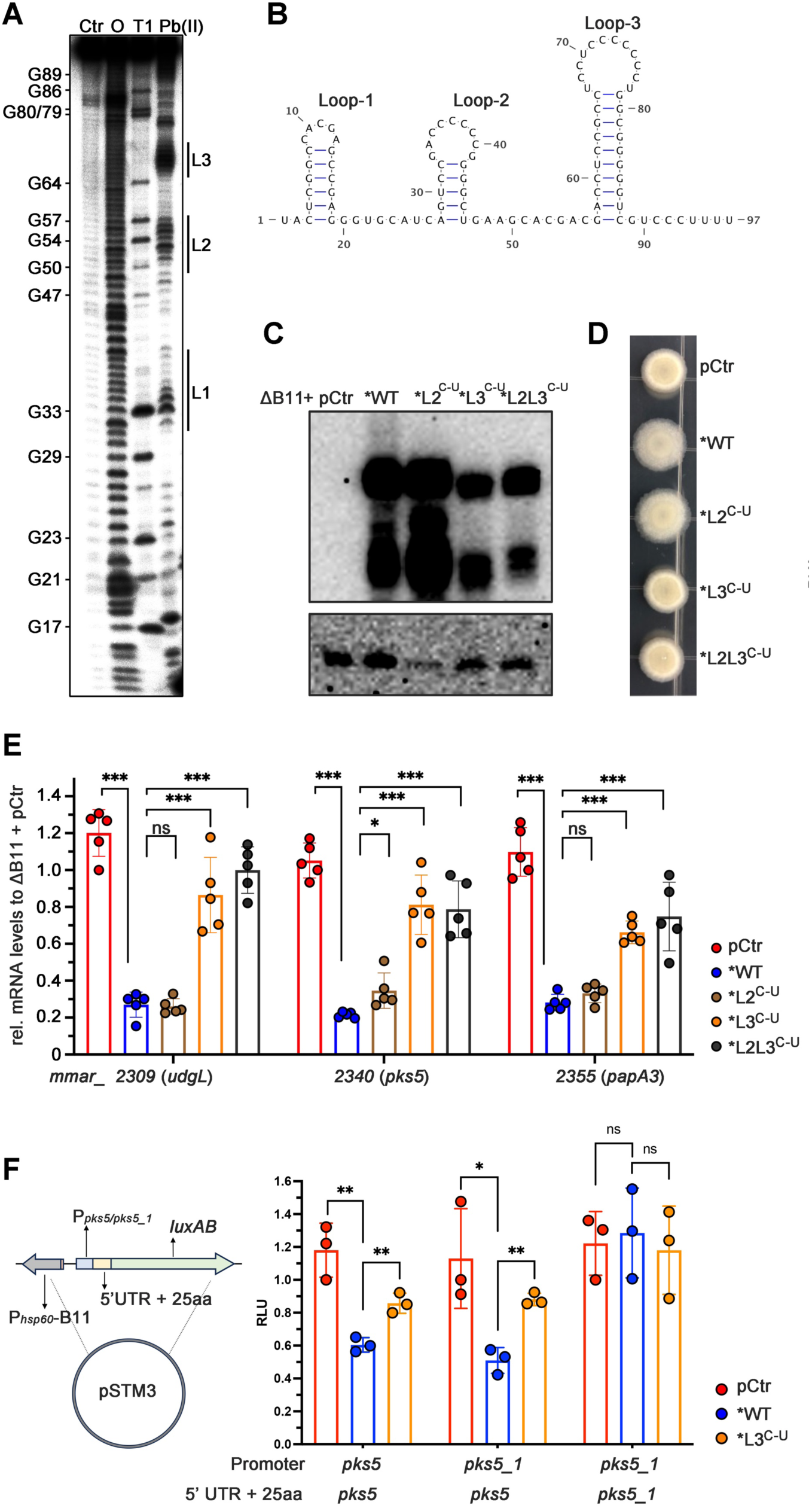
B11 regulates its targets through base pairing. (A) *In vitro* structure probing of B11 RNA using 5′ end-labelled B11 RNA with lead (II) acetate (PbII), RNase T1 (T1) and alkaline ladders (O) to map cleaved fragments. Positions of G-residues are indicated. (B) Illustrated secondary structure of B11 based on structure probing data. (C) Northern blot analysis of total RNA isolated from B11-deleted strain carrying different plasmids. 5S RNA was used as loading controls. (D) Morphology of indicated B11-deleted strain carrying different plasmids in 7H10 agar. (E) Expression of selected B11 targets in B11-deleted strain carrying different plasmids, confirmed by qRT-PCR. *sigA* was the reference gene for data analysis and expression levels were normalized to one sample from ΔB11 + pCtr group. The raw data from two groups (ΔB11 + pCtr and ΔB11 + pP*_b11_*-B11) are the same as shown in Figure 3B for comparison. Error bars indicate standard deviations (n = 5). (F) Relative RLU (relative light unit) of B11-deleted strain carrying plasmids containing the P*_MMAR_2340/2344_-*5’UTR of *mmar_2340/2344* -*luxAB* fusions. The results were normalized to one sample from the ΔB11 + pCtr group. Error bars indicate standard deviations (n = 3). * *p*<0.05, ** *p*<0.01, *** *p*<0.001, and *ns* refers to no significant difference, two-tailed *t* test.

To confirm that B11-mediated repression is post-transcriptional, we created a translation reporter system for mycobacteria based on the pXG-10 GFP reporter system in *E. coli* (39)(Figure 5F). B11 was expressed from the strong constitutive P*_hsp60_* promoter on plasmid pSMT3. On the same plasmid, the native promoters of the gene targets, along with their 5’ UTRs and first 25 amino acids, were fused to the second amino acid of *luxAB*. B11 and the *luxAB* fusion were expressed on different strands of the plasmid DNA to exclude the possibility of *cis*-acting regulation. To test the reporter system, we first chose *pks5* (*mmar_2340*), a B11 regulated gene encoding a type-I polyketide synthase. This gene has a homologue, *pks5_1* (*mmar_2344*), that is within in the same LOS biosynthetic locus but is not regulated by B11 and could therefore serve as a negative control. Bioluminescence assays showed that the presence of B11 in the WT strain reduced the relative luminescence units (RLU) of the *pks5 5’ UTR::luxAB* fusion nearly two fold (1.948 ± 0.149 fold) compared to the ΔB11 strain carrying the control plasmid without B11(Figure 5F), and mutation of the C-track in B11 loop-3 abolished this regulation. By contrast, no regulation was found for the *pks5_1 5’ UTR*::*luxAB.* To exclude the possibility of regulation at the transcriptional level, we replaced the *pks5* promoter in the fusion with the *pks5_1* promoter and found that the RLU was still reduced by 2.243 ± 0.071-fold, suggesting that it is the mRNA sequence and not the promoter that is regulated by B11. Taken together, our data strongly suggest that B11 regulates its targets at the post-transcriptional level, through the seed region located in the C-rich track of the 3’end loop 3.

### B11-mediated regulation is RNase E dependent

For most sRNAs in Gram-negative bacteria *E.coli* and *Salmonella*, base pairing around the mRNA RBS reduces ribosome binding frequency and leads to mRNA degradation, mainly by endoribonuclease RNase E (12). To determine whether B11 binding in *M. marinum* similarly leads to the degradation of target mRNAs, we evaluated B11 mediated-mRNA degradation in a strain in which RNase E (*rne*, *mmar_3768*) transcription was reduced by an anhydrotetracycline (ATc)-induced CRISPRi knock-down (40). At 24h after ATc induction, *rne* mRNA levels were reduced by ∼10 fold (Figure 6A), and concomitantly, the fold changes of targeted gene *udgL*, *pks5*, *mmar_2331* and *papA3* between cells carrying B11-overexpressed plasmid and empty control were decreased by 1.65 to 1.99-fold (Figure 6B) when *rne* expression was interfered, indicating that B11 promotes RNase E degradation of targeted mRNA.

**Figure 6.**
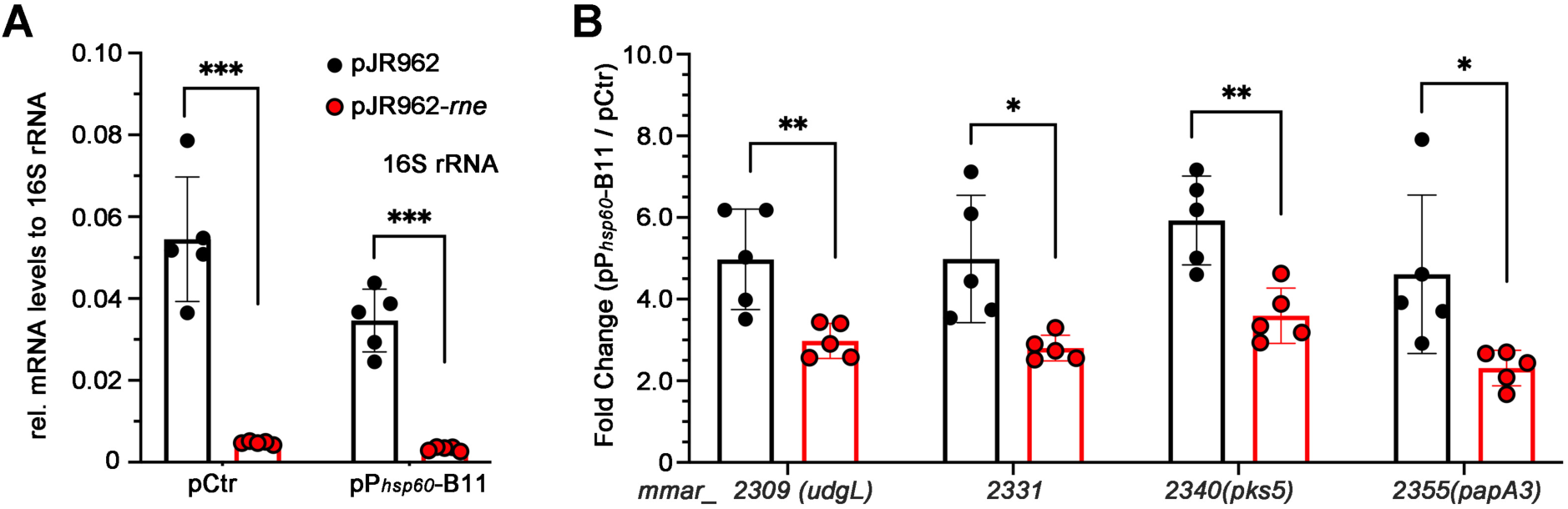
B11 mediated mRNA repression is RNase E dependent. (A) *rne* mRNA levels after 24h of exposure to ATc in the B11 deleted *M. marinum* strains containing either a control pSMT3 empty plasmid (pCtr) or B11 overexpressing plasmid (pP*_hsp60_*-B11) and also either the control plasmid pJR962 (black bars) or *rne-*interference plasmid pJR962-*rne* (red bars). 16S rRNA was set as the reference gene for data analysis. The vertical lines indicate standard deviations (n = 5). (B) Fold changes of B11-mediated repression (pP*_hsp60_*-B11 / pCtr) of 4 illustrative genes in strains with empty control plasmid pJR962 (black bars) or *rne-*interference plasmid pJR962-*rne* (red bars). Fold changes were calculated with normalized mRNA levels to 16S rRNA. Error bars indicate standard deviations (n = 5). * Indicates *p*<0.05. **, *p*<0.01. ***, *p*<0.00, and *ns*, no significant difference.

## Discussion

The extractable glycolipids of mycobacteria resemble the LPS of Gram-negative bacteria, both of which are characterized by genus conserved fatty acids (mycolic acid or lipid A) in the inner leaflets and species-specific lipids or glycans in the outer layer. Considering that LOS and LPS play key roles in adapting to varied environments and interacting with the host immune system, it is not surprising that their biosynthesis and assembly are controlled by multiple layers of regulation. In *E. coli* and many other Gram-negative bacteria, the regulation of LPS production is controlled both at the transcriptional level by transcriptional regulators and at the post-transcriptional level by sRNAs (41). For example, in *E. coli,* sRNA MgrR can fine-tune LPS structure by repressing the LPS-modification enzyme EptB(42). Another sRNA, MicF, regulates lipid A modification by promoting the degradation of *lpxR* mRNA, which encodes lipid A deacylase (43). In *V. cholerae*, sRNA VadR can regulates peptidoglycan integrity and cell shape by repressing expression of periplasmic protein CrvA(44). Although several conserved sRNAs have been identified in mycobacteria, only a few have been studied extensively for their biological roles and regulatory mechanisms, and none have been shown to be involved in cell wall synthesis. In this work we demonstrated sRNA-based regulation of cell wall synthesis in mycobacteria by performing a systematic identification of sRNA B11 targets. Through the use of several different approaches, including RNA sequencing and mass spectrometry screening, qPCR confirmation and MS2 affinity purification, we were able to identify genes involved in LOS synthesis that are regulated by B11. Although we did not analyze cell wall lipids to provide direct evidence of altered LOS production in the B11-deleted strain, its altered colony morphology is consistent with previously reported phenotypes in strains with deletions in LOS synthesis genes *udgL*(45,46), *pks5*(5), *fad25* (*mmar_2341*) and *papA3*(32). For example, inactivation of *udgL* (*mmar _2309*) and *mmar_2332* altered the colony morphology to exhibit wide translucent colony halos (46). This is consistent with our observation that in a B11-deleted strain these genes are up-regulated and the colonies have thin translucent halos, thereby confirming that B11 acts as a negative posttranscriptional regulator of LOS synthesis in *M. marinum*.

Because B11 is the most widely studied mycobacterial sRNA, it is possible to compare its function across different mycobacteria species, including *M. tuberculosis(19)*, *M. smegmatis* (26), *M. abscessus(27)*, and *M. kansasii*(47). However, the phenotype of B11-deficient strains has only been described in *M. abscessus*, *M. kansasii* and here, in *M. marinum*. Inactivation of B11 by transposon insertion in *M. kansasii* resulted in colonies with a reduced diameter, abnormal surface topology and a smooth and glistening appearance that contrasted with the rough and dried phenotype of WT colonies. Whether this mutant phenotype in *M. kansasii* is also caused by the loss of B11 repression of LOS biosynthesis is currently unknown, as the study did not identify B11 target genes (47). However, the genomic LOS biosynthesis locus in *M. kansasii*, particular the *pks5* region, resembles its orthologue in *M. marinum* (5). Furthermore, we identified a G-stretch close to the translation start site of the *M. kansasii pks5* gene *(mkan_27485,* an orthologue of *mmar_2340)*, suggesting that B11 regulation in *M. kansasii* could be similar to the B11 regulation we describe in *M. marinum*. In *M. abscessus,* which does not produce LOS, disruption of B11 resulted in smooth-to-rough colony variation by reducing the biosynthesis of glycopeptidolipids (GPL) through a mechanism, perhaps indirect, that appears to be independent of base-paring *(27)*. In addition, overexpression of B11 in the wild-type background of *M. tuberculosis* and *M. smegmatis* resulted in elongated bacilli, a phenotype that may be associated with the composition of the cell wall (19,26). Thus, although B11 appears to be involved with the regulation of cell wall synthesis in several mycobacteria species, the targeted genes and the regulatory mechanisms vary.

One surprising finding is that the direct B11 targets are distinct in the three species studied with target screening: *M. smegmatis* (26); *M. abscessus(27)*; and *M. marinum*. The direct B11 targets in *M. smegmatis* are involved with DNA replication and protein secretion, and in *M. abscessus* with ESX-related genes, but we found no evidence that B11 is involved with the regulation of these processes in *M. marinum*. Curiously, while it appears that the sequences of both the promoter and regulatory base-pairing seed regions of B11 are evolutionarily conserved in different *mycobacterium* species, and even in other *Corynebacterineae* families such as *Nocardiaceae* and *Gordoniaceae*, suggesting that the mechanism of regulation is also conserved, the biologic processes regulated are distinct in the different species. This raises the question: if B11 exhibits distinct behavior across various mycobacterium species, how does it maintain conservation throughout evolution? Studies on the conserved sRNA RyhB in *E. coli* and *Vibrio* suggest a model wherein broadly distributed sRNAs like RyhB, play a core role in iron homeostasis that is conserved across species(48,49). However, the *V. cholerae* sRNA has acquired additional physiological roles in both aerobic and anaerobic respiration(50,51). Drawing from this, we postulated that an unfolded core function of B11 may underpin its conservation among different mycobacteria. This function appears to deviate from the canonical mode of action through RNA-mRNA base pairing, potentially involving direct interaction with proteins. Further investigations are warranted to reveal this core function and ascertain whether B11 acts as a dual-function sRNA.

Our data showed that the seed region for regulatory base pairing by B11 is a cytosine-rich loop near its 3’ end, which appears to be conserved in the B11 orthologues in *M. smegmatis* (26) and *M. abscessus(27)*. Because mycobacteria have high GC-content with numerous guanine-rich motifs throughout the entire genome, computational methods using the cytosine-rich regions to predict the targets were not accurate. For instance, RNApredator predicted 5423 binding sites for B11(Table S6), and the locations of these sites covered almost all the annotated genes of *M. marinum*. Approximately 145 of these sites are predicted to have high stability (ΔG < -15). However, none of the genes associated with these 145 sites were identified in either our RNA sequencing or mass spectrometry data. In contrast, the primary target genes we identified, such as *udgL* and *mmar_4171*, were predicted to have relatively lower B11 binding (ΔG -12.01 and -10.55) and were not listed among the top candidates by RNApredator. Similarly, bioinformatic tools did not predict favorable binding energies for any of the experimentally confirmed targets of mycobacterial sRNA MrsI(25). The mode of sRNA-mRNA interaction in high GC-content mycobacteria may be different from that in other prokaryotes, perhaps requiring less favorable binding energy and independent of RNA chaperones such as Hfq and ProQ, which have not been found in mycobacterial genomes. Therefore, attempts to identify targets of mycobacterial sRNAs by bioinformatic scanning the genome for potential base-pairing sequences may be less fruitful than methods, such as those employed here, that can experimentally demonstrate sRNA-mRNA interactions. Methods similar like RIL-seq that are used in many bacteria (52-54), yet independent of capturing RNA-binding proteins, such as RIC-seq utilized in eukaryotic cells (55) are imperative for unraveling the rules governing sRNA-mRNA interactions in mycobacteria.

## Materials and Methods

### Bacterial strains and growth conditions

A complete list of bacterial strains used in this study is found in Table S1. *Mycobacterium marinum M* strain (ATCC BAA-535) was used throughout the study. Unless otherwise specified, strains were grown at 30°C in 7H9 medium (∼105 rpm) or 7H10 agar supplemented with 10% OADC and kanamycin (25 μg/ml), hygromycin B (20 μg/ml), or gentamycin (20 μg/ml) when necessary.

### DNA/RNA oligonucleotides and plasmids

All of the plasmids used in this study, with a brief description of their construction, are described in Table S2, and the sequences of the oligonucleotides used are listed in Table S3. Competent *E. coli* DH5α (#CB101, TIANGEN Biotech) were used for cloning. Plasmids were isolated using the TIANprep Rapid Mini Plasmid Kit (#DP105, TIANGEN Biotech) and confirmed by Sanger sequencing (Shanghai Personal Biotechnology Co., Ltd).

### Sequence alignments

Nucleotide Blast searches (https://blast.ncbi.nlm.nih.gov/Blast.cgi) were performed with the following genomes: *Mycobacterium marinum M* (NC_010612.1); *Mycobacterium ulcerans Agy99* (NC_008611.1); *Mycobacterium kansasii* ATCC 12478 (NC_022663.1); *Mycobacterium tuberculosis H37Rv* (NC_000962.3); *Mycobacterium tuberculosis H37Ra* (NC_009525.1); *Mycobacterium canettii CIPT 140070007* isolate STB-I (NZ_CAOO00000000.1); *Mycobacterium bovis BCG str. Tokyo* (AP010918.1); *Mycobacterium africanum K85* (KK338483.1); *Mycobacterium avium subsp. paratuberculosis K-10* (NC_002944.2); *Mycobacterium intracellulare ATCC 13950* (NC_016946.1); *Mycobacterium indicus pranii MTCC 9506* (CP002275.1); *Mycobacterium smegmatis str. MC2 155* (NC_008596.1); *Mycolicibacterium vanbaalenii PYR-1* (NC_008726.1); *Mycolicibacterium gilvum Spyr1* (NC_014814.1); Nucleotide sequence alignments were generated with MultAlin http://multalin.toulouse.inra.fr/multalin/multalin.html.

### Construction of ΔB11 strain

Deletion of B11 in *M.marinum* was achieved by a modified allelic exchange method using the temperature-sensitive pPR27 plasmid(31). This vector harbors a thermosensitive origin of replication and the *sacB* gene from *B. subtilis*, which is lethal to mycobacteria in the presence of sucrose. However, the temperature generally used for losing the plasmid, 39 °C, is not suitable for *M. marinum* growth. To solve this, we inserted the *mWasabi* gene to create a pPR27-*mWasabi* plasmid that expresses a brighter green fluorescent protein. *M. marinum* strains carrying this plasmid have a green color that distinguishes them from white strains that have lost the plasmid, which made it possible to perform the selection at 30°C. For allelic exchange, 1kb downstream of b11 nucleotides 36-91 was amplified by PCR with oligo QGO-037/038, and 1kb upstream was amplified with oligo QGO-039/040. The two fragments were fused on either side of a kanamycin resistance gene amplified from pMV306 plasmid with oligos QGO-041/042, to yield a ∼3.2 kb fragment with the kanamycin resistance gene in the middle. The fused product was inserted into pPR27-*mWasabi* and transformed into the wild-type *M. marinum M* strain. After 10 days of growth, kanamycin and gentamicin double resistant, GFP-positive colonies was selected from 7H10 plates and incubated in 7H9-OADC media supplemented with kanamycin and gentamicin to OD600∼0.5. The bacterial culture was collected and filtered through a 5 μM filter (Sartorius) to obtain a single-bacteria cell suspension. The suspension was then serially diluted and plated on 7H10-agar containing 10% sucrose to inhibit growth of cells retaining the pPR27 plasmid. After 20 days, one colorless clone was found among ∼5000 green colonies on 100 plates. After confirming the loss of fluorescent with the Typhoon FLA 9000 system (GE), the colorless clone was incubated in 7H9-OADC supplemented with kanamycin to OD600∼0.5 and then streaked on 7H10-OADC plates to isolate a pure strain whose B11 deletion was confirmed by PCR and northern blotting.

### RNA isolation and Northern blot analysis

4 OD_600_ bacterial cultures were mixed with 0.2 volumes of stop solution (95% ethanol and 5% phenol) and immediately frozen in liquid nitrogen. Total RNA was isolated using the Trizol method. Briefly, cells were resuspended with 1mL Trizol (#15596018, Invitrogen) and 0.2 mL 0.1mm Zirconium/Silica beads (#11079101z, Biospec). Bacteria were lysed with a Precellys 24 homogenizer (4500 rpm, 3x30s, separated by 5min intervals on ice,). After mixing with 400 μl chloroform and centrifugation in a Phase Lock Gel tube (#WM5-2302820, TIANGEN Biotech) the samples were centrifuged at 13,000 rpm for 15 min, and then 500 ul of the aqueous phase was mixed with 450 μl isopropanol and precipitated at room temperature for 30 min. After centrifugation at 13,000 rpm for 30min, the RNA pellets were washed with 80% ethanol, dissolved in ultra-pure water (#10977, Thermo Fisher Scientific), and the RNA concentration was determined with a NanoDrop 2000 (Thermo Fisher Scientific).

For northern blot analysis, 10 μg of total RNA was denatured at 95°C for 2 min in RNA gel loading buffer II (95% formamide, 18 mM EDTA, 0.025% SDS, 0.025% xylene cyanol, 0.025% bromophenol blue), and separated by gel electrophoresis on 6% polyacrylamide/7 M urea gels in 1x TBE buffer for 2h at 300 V using the Beijing Liuyi DYCZ system (Beijing Liuyi Biotechnology). RNA was transferred from gels onto Hybond-N+ membranes (GE Healthcare) by electroblotting for 2 h at 25 V at 4°C. The membranes were prehybridized in DIG Easy Hyb (#11796895001, Roche) for 30 min and hybridized with the Digoxin-labelled DNA probe at 50°C overnight. The membranes were then washed for 15 minutes each in 5× SSC/0.1% SDS, 1× SSC/0.1 % SDS and 0.5x SSC/0.1 % SDS at 50°C. After incubations in maleic acid wash buffer (0.1 M maleic acid, 0.15 M NaCl, 0.3% Tween-20 pH 7.5) for 5 min at 37°C and blocking solution (#11585762001, Roche) for 45 min at 37°C, the membrane was incubated with 75 mU/mL Anti-Digoxigenin-AP (#11093274001, Roche) in blocking solution for 45 min at 37°C. Following two 15-minute maleic acid washes, the membrane was equilibrated with detection buffer (0.1 M Tris-HCl, 0.1 M NaCl, pH 9.5) and then signals were visualized by CDP-star (#12041677001, Roche) on a ChemiDocTM XRS+ station and quantified using ImageLabTM software (Biorad).

### RNA sequencing

RNA-seq was performed by the BGI Group, Shenzhen, Guangdong, China. Briefly, a non-strand-specific cDNA library was constructed for each sample. The cDNA libraries were then sequenced with a BGISeq-500 instrument. Adaptors were removed and the quality of RNA-seq data was assessed with the SOAP software package(56). High-quality reads were mapped to the genome of *Mycobacterium marinum* M (NC_010612.1) using the HISAT software package(57). Relative expression of each individual gene was calculated in each sample by counting fragments per kilobase of transcript per million mapped reads (FPKM) with Bowtie2 and RSEM software packages(58,59).

### Mass spectrometry

The mass spectrometry proteomics analysis was performed by Shanghai iProteome Biotechnology Co., Ltd. Briefly, ∼50 OD_600_ bacteria cells cultivated in 7H9-OADC medium were collected and washed three times with phosphate buffer saline (PBS) buffer. Samples were lysed in lysis buffer (8 M urea, 100 mM Tris hydrochloride, pH 8.0) containing protease and phosphatase inhibitors and then sonicated for 1 min (3 s on and 3 s off, amplitude 25%). The lysates were centrifuged at 14,000 × g for 10 min and supernatants were collected. Protein concentrations were determined by the Bradford protein assay. Extracts (1mg protein) were reduced with 10 mM dithiothreitol at 56 °C for 30 min and alkylated with 10 mM iodoacetamide at room temperature in the dark for 30 min. The samples were digested with trypsin using a filter-aided sample preparation method84. Tryptic peptides were vacuum-dried (Concentrator Plus, Eppendorf). For the proteome profiling samples, peptides were analyzed on a Q-Exactive HFX Hybrid Quadrupole-Orbitrap Mass Spectrometer (Thermo Fisher Scientific, Rockford, IL, USA) coupled with a high-performance liquid chromatography system (EASY nLC 1200, Thermo Fisher). The original data of mass spectrometry analysis were. d files, and MaxQuant was used for qualitative and quantitative analysis. The fraction of total (FOT), a relative quantification value was defined as a protein’s intensity-based absolute quantification (60).

### Quantitative PCR

Quantitative PCR was performed with the PrimeScript™ RT reagent Kit (#RR047A, Takara Bio). Briefly, 1 μg total RNA was treated with gDNA eraser, provided in the kit, at 42°C for 2 min, and then reverse-transcribed to cDNA with random oligos and PrimeScript RT Enzyme Mix I at 37°C for 15 min. cDNA transcribed from 0.025 μg total RNA was used per PCR reaction in a final volume of 20 μl with TB Green Premix Ex Taq II (#RR820A, Takara Bio). PCR was performed using a Biorad CFX96 Touch Real-Time PCR Detection System. Data were analyzed by the relative quantification (ddCt) method, with the rfaH gene as the reference for normalization.

### *In vitro* RNA synthesis and RNA labelling

For *in vitro* synthesis of B11 transcripts, 200 ng of a DNA fragment amplified from M. marinum genomic DNA with primer pair QGO-782/784 served as the template in a T7 transcription reaction using the MEGAscript™ T7 Transcription Kit (#AM1334, Thermo Fisher Scientific). The correct size and integrity of RNA were confirmed in a denaturing polyacrylamide gel. RNA was excised from the gel and eluted in 0.1 M sodium acetate, 0.1 % SDS, 10 mM EDTA at 4°C overnight, followed by phenol:chloroform:isoamyl extraction and precipitation. For radio-labelling, 50 pmol RNA was dephosphorylated with 10 units of calf intestine alkaline phosphatase (#M0290, New England Biolabs) in a 50 μl reaction at 37°C for 1 h, followed by phenol:chloroform:isoamyl extraction and precipitation. 20 pmol dephosphorylated RNA was 5′-labelled with 3 μl of 32P-γ-ATP (10 Ci/l, 3,000 Ci/mmol) using 1 unit of T4 polynucleotide kinase (#EK0031, Thermo Fisher Scientific) for 1 h at 37°C in a 20 μl reaction. Microspin G-50 columns (#27533001, GE Healthcare) were used to remove unincorporated nucleotides, according to the manufacturer’s instructions.

### RNA structure probing

Structure probing was performed with in vitro transcribed 5′-radio-labelled RNA as described(17). Briefly, 0.2 pmol radio-labelled RNA in a 5 μl volume was denatured at 95°C for 2 min and chilled on ice for 5 min, followed by the addition of 1 μl of yeast RNA (1 mg/ml, #AM7118, Thermo Fisher Scientific), and 1 μl of 10× structure buffer (0.1 M Tris– HCl, pH 7, 1 M KCl, 0.1 M MgCl2) and then incubation at 37°C for 30 min. Samples were treated with 5 mM lead (II) acetate (Fluka) for 1.5 min. RNase T1 sequencing ladders were prepared using 0.4 pmol of 5′-labelled RNA digested with 0.1 U RNase T1 for 5 min at 37°C. Alkaline sequencing ladders were prepared by incubating 0.4 pmol of 5′-labelled RNA at 95°C for 5 min in the presence of alkaline hydrolysis buffer. Reactions were stopped by adding 12 μl RNA Gel loading buffer II. Samples were denatured at 95°C for 2 min and separated on 8 % denaturing sequencing gels containing 7 M urea in 1× TBE at constant power of 40 W for 1 h. Gels were dried and signals were analyzed on a Typhoon FLA 7000 phosphoimager using AIDA software.

### MS2 affinity purification

MS2 affinity purification was performed as described(36). MS2-B11 expressed from the Hsp60 promoter showed expression and regulatory ability similar to B11 expressed from its native promoter, the ΔB11 + pPb11-B11 strain was used as the control for the affinity purification. Briefly, 50 OD_600_ bacterial cultures at OD_600_∼1.0 were washed and resuspended in 10 mL ice-cold buffer A (20 mM Tris–HCl at pH 8.0, 150mM KCl, 1mM MgCl2, 1mM DTT, and 1mM PMSF.) and frozen immediately in liquid nitrogen. These samples were mixed with 0.1mm Zirconia/Silica beads (Biospec) and disrupted with a Precellys 24 homogenizer (6800 rpm, 5x30seconds, separated by 10 min intervals in liquid nitrogen). The lysates were cleared by centrifugation at 15,000 rpm, 4°C for 30 min and supernatants were collected. Before affinity purification, 75 μL amylose resin (#E8021S, New England Biolabs) was added to a Bio-Spin disposable chromatography column (#7326008, Bio-rad) and equilibrated three times with 1 mL Buffer A. 100 pmol purified MS2-MBP coat protein was then diluted in 1 mL Buffer A and applied onto the column. After 5 min incubation and two column washes with Buffer A, 10mL of bacteria lysate was loaded onto the column. The column was washed five times with 1 mL Buffer A and then eluted with 1 mL Buffer A supplemented with 15 mM maltose. Eluted RNA was isolated by phenol:chloroform:isoamyl extraction and precipitation as described for RNA labelling. 50 ng RNA, quantitated with a 5300 Fragment Analyzer System (Agilent) was then used for qRT-PCR reactions.

### Bioluminescence assay

Bacterial cultures at OD_600_∼1.0 were serially diluted and plated on 7H10 -OADC agar. For the bioluminescence assay, 0.1 mL culture was added into the wells of white 96-well plates (#3917, Corning). 1% decanal (#D7384, Sigma-aldrich) in ethanol was 1:25 diluted with 7H9 medium to a final concentration of 0.04% to make the reaction buffer. Subsequently, 0.1 mL reaction buffer was added to the wells with bacterial cultures and bioluminescence was detected by a FlexStation 3 Multi-Mode Microplate Reader (Molecular Devices).

### Statistical analyses

Statistical parameters for each experiment are detailed in the respective figure legends. ‘n’ denotes the number of biological replicates. Statistical analysis utilized GraphPad Prism 9 (GraphPad Software). Experiments were not blinded or randomized, and no prior estimation of statistical power was conducted. Additionally, no data were excluded from analysis.

## DATA AVAILABILITY

The data supporting the findings of this study are available from the corresponding authors upon request. The RNA sequencing data have been deposited in the GEO database under No. GSE264453. Mass spectrometry data have been deposited in the iProX database under PXD051635.

## ACKNOWLEDGEMENTS

We thank Yanjie Chao for comments on the manuscript. This work was supported by the National Key R&D Program of China (2022YFE0111800 to C.W.), the Natural Science Foundation of China (82272376 to Q.G. 81991532 and 32170179 to C.W) and Shanghai Municipal Science and Technology Major Project (ZD2021CY001).

## CONFLICT OF INTEREST

The authors declare that they have no conflict of interest.

## Author contributions

C.W., and Q.G. designed the experiments; C.W., Y.F., Y.D. performed the experiments; C.W., C.B., and Q. L. analyzed data; and C.W. wrote the manuscript with the help of all authors.

